# Cardiovascular effects of intravenous colforsin in normal and acute respiratory acidosis canine models: a dose–response study

**DOI:** 10.1101/558221

**Authors:** Takaharu Itami, Kiwamu Hanazono, Norihiko Oyama, Tadashi Sano, Kazuto Yamashita

**Author notes:** ^*^Corresponding author Takaharu Itami, D.V.M., Ph.D. 582, Bunkyodai-Midorimachi, Ebetsu, Hokkaido 069-8501, Japan, E-mail. (TI). These authors contributed equally to this work (Takaharu Itami, Kiwamu Hanazono).

## Abstract

**Abstract:** In acidosis, catecholamines are attenuated and higher doses are often required to improve cardiovascular function. Colforsin activates adenylate cyclase in cardiomyocytes without mediating the beta adrenoceptor. In this study, six beagles were administered either colforsin or dobutamine four times during eucapnia (partial pressure of arterial carbon dioxide 35-40 mm Hg; normal) and hypercapnia (ibid 90-110 mm Hg; acidosis) conditions. The latter was induced by carbon dioxide inhalation. Anesthesia was induced with propofol and maintained with isoflurane. Cardiovascular function was measured by thermodilution and a Swan-Ganz catheter. Cardiac output, heart rate, and systemic vascular resistance were determined at baseline and 60 min after 0.3 μg/kg/min (low), 0.6 μg/kg/min (middle), and 1.2 μg/kg/min (high) colforsin administration. The median pH was 7.38 [range 7.34–7.42] and 7.04 [range 7.01–7.08] at baseline in the Normal and Acidosis conditions, respectively. Endogenous adrenaline and noradrenaline levels at baseline were significantly (*P* < 0.05) higher in the Acidosis than in the Normal condition. Colforsin induced cardiovascular effects similar to those caused by dobutamine. Colforsin increased cardiac output in the Normal condition (baseline: 198.8 mL/kg/min [range 119.6–240.9], low: 210.8 mL/kg/min [range 171.9–362.6], middle: 313.8 mL/kg/min [range 231.2–473.2], high: 441.4 mL/kg/min [range 373.9–509.3]; *P* < 0.001) and the Acidosis condition (baseline: 285.0 mL/kg/min [range 195.9–355.0], low: 297.4 mL/kg/min [213.3–340.6], middle: 336.3 mL/kg/min [291.3–414.5], high: 366.7 mL/kg/min [339.7–455.7] ml/kg/min; *P* < 0.001). Colforsin significantly increased heart rate (*P* < 0.05 in both conditions) and decreased systemic vascular resistance (*P* < 0.05 in both conditions) compared to values at baseline. Systemic vascular resistance was lower in the Acidosis than in the Normal condition (*P* < 0.001). Dobutamine increased pulmonary artery pressure, whereas colforsin did not. Colforsin offsets the effects of endogenous catecholamines and may not increase cardiac output during hypercapnia.

## Introduction

Catecholamine beta-1 adrenoceptor is present on myocardial cell membranes. The catecholamine dobutamine binds to the beta-1 adrenoceptor and activates the cyclic adenosine monophosphate (cAMP) synthetase adenylate cyclase. The cAMP activates protein kinase A which phosphorylates the L-type calcium ion- and sarcoplasmic reticulum calcium ion-releasing channels and increases intracellular calcium ion concentrations. Dobutamine increases cardiac contractility and heart rate [1]. In contrast, catecholamine beta-2 adrenoceptor is present on the vascular smooth muscle cell membrane. Protein kinase A activated as described above phosphorylates myosin light-chain kinase and inhibits actin and myosin gliding. Dobutamine relaxes vascular smooth muscle, has both positive inotropic and vasodilator effects (inodilator), and is cardiotonic and vasodilatory action in dogs [2].

Colforsin daropate is a forskolin derivative that directly activates adenylate cyclase in cardiomyocytes and vascular smooth muscle without mediating the catecholamine beta adrenoceptor. As with dobutamine, colforsin increases cardiac contractility and reduces peripheral vascular resistance [3,4]. Colforsin has been tested on human patients with congestive heart failure and it improved their hemodynamics [5]. When forskolin was first discovered, it was poorly soluble in water and its clinical application as an injection was limited. Colforsin was prepared as a water-soluble forskolin derivative and became available in 1999 [5]. However, the efficacy of colforsin was never compared with that of catecholamines and it was not tested on animals or humans until now. Catecholamines and phosphodiesterase III inhibitors have been reported as inodilators in dogs. To the best of our knowledge, however, there have been no reports on the cardiovascular effects of colforsin in dogs.

In sepsis and acidosis, myocardial beta-1 adrenoceptor is downregulated. Therefore, catecholamine responses to decreases in cAMP decline. For this reason, cardiac contractility is suppressed in sepsis and acidosis [6]. Unlike adrenaline, colforsin improved cardiac function in rat cardiac resection specimens even under acidosis [7]. However, no investigation has been conducted on the effects of colforsin in living organisms under acidosis. We hypothesized that colforsin maintains cardiac contractility in acidotic dogs.

The purposes of this study were to: 1) investigate the cardiovascular effects of colforsin in dogs and 2) examine the cardiovascular effects of colforsin in an acute respiratory acidosis model induced by carbon dioxide inhalation.

## Materials and methods

### Experimental animals

Six beagles (3 females, 3 males) aged 1–2 y and weighing 9.5–12.5 kg (mean ± SD: 10.9 ± 1.0 kg) were used in this study. The dogs were judged to be in good health based on the results of physical examinations, complete blood cell counts, and serum biochemical analyses. The dogs were owned by the university and maintained according to the principles of the “Guide for the Care and Use of Laboratory Animals” prepared by Hokkaido University and approved by the Association for Assessment and Accreditation of Laboratory Animal Care International (AAALAC). The Animal Care and Use Committee of Hokkaido University approved the study (No. 14-0156). Food (but not water) was withheld from the dogs for 12 h before the experiment. Dogs in normal and acidotic condition were administered colforsin or dobutamine. Each dog was anesthetized 4× at 2-week intervals. This study was performed in a randomized crossover design.

### Experimental preparations

All dogs were fitted with 22-gauge catheters (Surflow; Terumo Co. Ltd., Tokyo, Japan) in both cephalic veins and administered 6 mg/kg propofol (Propoflo 28; Zoetis Co. Ltd., Tokyo, Japan) intravenously through a catheter placed in the right cephalic vein. They were orotracheally intubated and connected to a standard circle anesthesia system (FO-20A; ACOMA Medical Industry Co. Ltd., Tokyo Japan) and a ventilator (Spiritus; ACOMA Medical Industry Co. Ltd., Tokyo Japan). All dogs received Ringer’s solution (Fuso Pharmaceutical Industries Ltd., Osaka, Japan) at 5 mL/kg/h and vecuronium (Fuji Pharma Co. Ltd., Tokyo, Japan) by injection at 0.1 mg/kg. They were then intravenously infused with 0.1 mg/kg/h vecuronium through a catheter in the right cephalic vein to prevent reflex respiratory muscle movement. The dogs were mechanically ventilated with oxygen and received 1.3-1.5% end tidal isoflurane anesthesia. The oxygen flow rate was 2 L/min. They were placed in left lateral recumbency and mechanically ventilated at a respiratory rate of 12 breaths/min and a 1:2 inspiratory-expiratory ratio with volume control ventilation (tidal volume = 9–13 mL/kg).

A 22-gauge catheter was inserted percutaneously into a left dorsal pedal artery. Three pressure transducers (DT-NN; Merit Medical Co. Ltd., Tokyo, Japan) were prepared and calibrated against a mercury manometer at 200 mm Hg, 50 mm Hg, and 20 mm Hg for the mean arterial, pulmonary arterial, and right atrial pressures, respectively. The right neck region was shaved and aseptically prepared. Approximately 0.5 mL of 2% lidocaine (xylocaine; Astra-Zeneca, Osaka, Japan) was injected subcutaneously. A 5-Fr, 75-cm Swan-Ganz catheter (132F5; Edwards Lifesciences Co. Ltd., Tokyo, Japan) was inserted into a jugular vein using a 6-Fr introducer (Medikit Catheter Introducer; Medikit Co. Ltd., Tokyo, Japan). The distal port of the Swan-Ganz catheter was connected to a pressure transducer and advanced into the pulmonary artery using the characteristic pressure changes associated with the right ventricle and pulmonary artery. A transducer was attached to the arterial catheter to measure mean arterial pressures (MAP; mm Hg). Transducers were connected to the distal and proximal ports of the Swan-Ganz catheter to measure mean pulmonary arterial pressure (PAP; mm Hg) at the distal port, pulmonary arterial occlusion pressure (PAOP; mm Hg) at the distal port, and mean right atrial pressure (RAP; mm Hg) at the proximal port. All pressure transducers were zeroed at the mid-sternum level. The PAOP was measured after distal balloon inflation on the Swan-Ganz catheter at the end of expiration.

Cardiac output (CO; L/min) was determined by thermodilution. Five milliliters of normal saline (1-4 °C) was rapidly injected manually into the proximal port of the Swan-Ganz catheter at the end of expiration. Temperature fluctuations were measured with a thermosensor placed at the tip of the Swan-Ganz catheter. At each time interval, three consecutive measurements within 10% of each other were recorded and the average was recorded as the CO. The thermistor on the Swan-Ganz catheter measured the core body temperature which was maintained between 37.0–37.5 °C by a forced-air patient-warming machine (Bair Hugger; 3M Japan Co. Ltd., Tokyo, Japan).

After the dogs were instrumented, the normal and acidotic conditions were adjusted according to the arterial blood gas data. Arterial- and mixed venous blood gases were measured by collecting 1.0 mL blood from the dorsal pedal- and pulmonary arteries catheterized to a heparinized syringe. Blood gas measurements (GEM-Premier 3000; IL Japan Co. Ltd., Tokyo, Japan) were corrected to body temperature. When the cardiovascular parameters were being measured in the normal condition (Normal), the arterial partial pressure of carbon dioxide (PaCO_2_) was maintained at ~35–40 mm Hg. When the cardiovascular parameters were being measured in the acidotic condition (Acidosis), the PaCO_2_ was maintained at ~90–110 mm Hg and the pH was ~7.0. Exogenous hypercapnia was induced by adding dry gaseous carbon dioxide (CO_2_) to the inspiratory corrugated tube of the anesthesia circuit.

### Evaluation of cardiovascular parameters

All dogs were stabilized for 30 min after preparation. Then, baseline cardiovascular parameters and arterial- and mixed venous blood gases were measured and recorded as follows: heart rate (HR; beats/min) by electrocardiogram with a lead II, and MAP, PAP, RAP, and PAOP by a multi-parameter anesthetic monitoring system (RMC-4000 Cardio Master; Nihon Kohden Corporation, Tokyo, Japan). Cardiac index (CI; mL/min/kg), stroke volume (SVI; mL/beat/kg), systemic vascular resistance (SVRI; dynes·sec·cm^−5^/kg), and pulmonary vascular resistance (PVRI; dynes·sec·cm^−5^/kg) were calculated by inserting values into previously published formulae [8].

After the baseline measurements, the dogs were intravenously infused with colforsin (Adehl; Nihonkayaku Co. Ltd., Tokyo, Japan) or dobutamine (Dobutrex; Shionogi & Co. Ltd., Osaka, Japan) through a 22-gauge catheter inserted into the left cephalic vein. Colforsin administration was gradually increased to 1 mL/h (0.3 μg/kg/min), 2 mL/h (0.6 μg/kg/min), and 4 mL/h (1.2 μg/kg/min) every 60 min. Similarly, dobutamine administration was gradually increased to 1 mL/h (5 μg/kg/min), 2 mL/h (10 μg/kg/min), and 4 mL/h (20 μg/kg/min) every 60 min. Colforsin and dobutamine were diluted with sterile saline (normal saline; Otsuka Pharmaceutical Factory Inc., Tokyo, Japan) and administered by infusion pump (TOP-5500; TOP Co. Ltd., Tokyo, Japan). All cardiopulmonary measurements were repeated every 60 min after each dose was administered. When the cardiovascular parameters were determined after the final dose, the arterial- and mixed venous blood gases were measured as described above.

After the experiment, all dogs received 0.2 mg/kg subcutaneous meloxicam (Metacam; Boehringer Ingelheim Co. Ltd., Tokyo, Japan) and 0.01 mg/kg intramuscular buprenorphine (Lepetan injection; Otsuka Pharmaceutical Factory Inc., Tokyo, Japan) for analgesia and 25 mg/kg intravenous cefazolin (cefazolin sodium; Nichi-Iko Co. Ltd., Toyama, Japan) to prevent infection. For the Acidosis condition, carbon dioxide inhalation was terminated and dog PaCO_2_ was maintained at 35–40 mm Hg. They were administered 0.5 g/kg intravenous mannitol (*D*-mannitol injection; Terumo Co. Ltd., Tokyo, Japan) for 30 min to lower intracranial pressure. Colforsin and dobutamine were washed out for 1 h and the dogs recovered from the anesthesia.

### Biochemical examination

Two milliliters of blood was drawn from the arterial catheter to measure baseline catecholamine (adrenaline, noradrenaline, and dopamine) concentrations. The blood samples were immediately centrifuged (1,000 × *g* for 10 min at 4° C) to separate the plasma, which was then stored at −80 °C until analysis. Catecholamine levels were determined by an external laboratory (BML Inc., Tokyo, Japan). In addition, 2 mL blood was drawn from the arterial catheter at baseline and at the end of experiment, the plasma was isolated from them as described above, and the samples were biochemically analyzed (DRI-CHEM 7000V; Fujifilm Co. Ltd., Tokyo, Japan) in the laboratory at our facility.

### Statistical analysis

The data were processed using a statistical software (BellCurve for Excel; Social Survey Research Information Co. Ltd., Tokyo, Japan) and online in R v. 3.5.0. (2018-04-23). A Wilcoxson signed-rank test was used to compare biochemical measurements between baseline and the end of the experiment. It was also used to compare cardiovascular variables at each colforsin and dobutamine dose between the Normal and Acidosis conditions. A rank transformation version of two-way ANOVA was used to compare the Normal and Acidosis conditions. A post-hoc Steel-Dwass test was used to compare dose-related effects on cardiovascular parameters. Differences were considered significant when *P* < 0.05.

## Results

### Blood gases and biochemical analyses in response to colforsin

The blood gas and biochemical test results at baseline and at the end of the experiment are shown in Table 1. The median pH at baseline was 7.38 (range 7.34–7.42) for the Normal condition and 7.04 (7.01–7.08) for the Acidosis condition. In both cases, the pH was slightly lower by the end of the experiment. The PaCO_2_ at baseline was 39.5 mm Hg (range 34.0–41.1 mm Hg) for the Normal condition and 97.8 mm Hg (range 92.0–100.4 mm Hg) for the Acidosis condition.

**Table 1.**
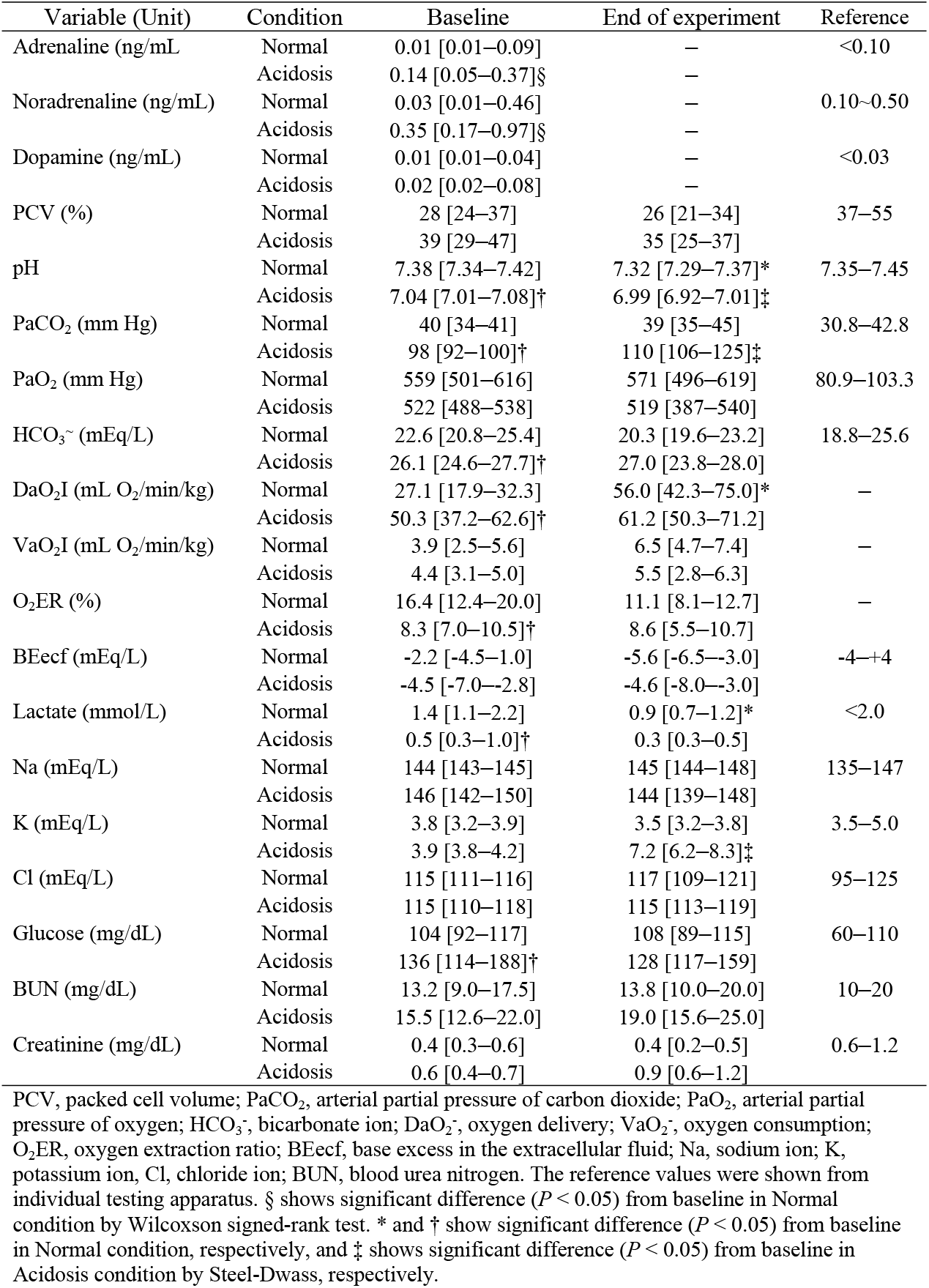
The effects of colforsin on blood gas examination and blood biochemical test in six anesthetized dogs in eucapnia (Normal) and acute respiratory acidosis (Acidosis) at baseline and at the end of the experiment.

Blood adrenaline and noradrenaline levels were significantly higher in the Acidosis than in the Normal condition at baseline (*P* < 0.05 for both). Plasma glucose level was significantly (*P* < 0.05) higher in the Acidosis condition than the Normal condition. In the former, plasma potassium was significantly (*P* < 0.05) higher at the end of the experiment than it was at baseline. Although the peak plasma potassium was 8.3 mmol/L, no arrhythmia was observed.

### Cardiovascular effects of colforsin

The effects of colforsin on the cardiovascular parameters in the Normal and Acidosis conditions are shown in Table 2. There was an interaction between the effects of colforsin and pH on both CI and SVRI. Baseline CI, HR, SVI, PAP, and PAOP were higher and SVRI was lower in the Acidosis condition than the Normal condition, and the differences were significant (*P* < 0.05). Relative to the baseline value, the rate of increase in CI in the Acidosis condition was greater than that in the Normal condition (Normal vs. Acidosis: 6% vs. 4%, 0.3 μg/kg/min; 58% vs. 18%, 0.6 μg/kg/min; 122% vs. 29%, 1.2 μg/kg/min). The SVI, DAP, RAP, SVRI, and PAOP were significantly (*P* < 0.05) different between the Normal and Acidosis conditions.

**Table 2.** Median [range] values for cardiovascular variables at baseline, 0.3 μg/kg/min, 0.6 μg/kg/min, and 1.2 μg/kg/min dose of intravenous colforsin in six anesthetized dogs in eucapnia (Normal) and acute respiratory acidosis (Acidosis) conditions.

| Variable (Unit) | Condition | Baseline | 0.3 µg/kg/min | 0.6 µg/kg/min | 1.2 µg/kg/min | P-value |  |  |
| --- | --- | --- | --- | --- | --- | --- | --- | --- |
|  |  |  |  |  |  | Condition | Treatment | Interaction |
| CI (mL/min/kg) | Normal | 198.8 [119.6–240.9] | 210.8 [171.9–362.6] | 313.8 [231.2–473.2]<sup>A</sup> | 441.4 [373.9–509.3]<sup>AB</sup> | 0.189 | <0.001 | 0.033 |
|  | Acidosis | 285.0 [195.9–355.0]* | 297.4 [213.3–340.6] | 336.3 [291.3–414.5] | 366.7 [339.7–455.7]*<sup>B</sup> |  |  |  |
| HR (beats/min) | Normal | 81 [68–116] | 99 [76–140] | 152 [95–173] | 197 [188–209]<sup>ABC</sup> | 0.194 | <0.001 | 0.071 |
|  | Acidosis | 114 [94–133]* | 120 [99–131] | 129 [107–155] | 135 [123–165]* |  |  |  |
| SVI (mL/beat/kg) | Normal | 1.86 [1.46–2.65] | 2.13 [2.03–2.57] | 2.37 [1.88–2.63] | 2.06 [1.88–2.34] | <0.001 | 0.106 | 0.356 |
|  | Acidosis | 2.51 [1.97–2.67]* | 2.59 [2.16–2.75]* | 2.64 [2.41–2.74] | 2.68 [2.53–2.89]* |  |  |  |
| SAP (mm Hg) | Normal | 101 [68–114] | 101 [82–115] | 88 [84–110] | 81 [70–93] | 0.176 | 0.014 | 0.894 |
|  | Acidosis | 96 [81–98] | 87 [85–100] | 92 [61–102] | 79 [49–101] |  |  |  |
| MAP (mm Hg) | Normal | 71 [50–84] | 74 [56–86] | 66 [58–77] | 59 [36–66] | 0.335 | 0.033 | 0.835 |
|  | Acidosis | 69 [51–77] | 67 [55–76] | 65 [42–74] | 57 [36–76] |  |  |  |
| DAP (mm Hg) | Normal | 57 [40–67] | 60 [40–75] | 58 [44–60] | 50 [42–64] | <0.001 | 0.078 | 0.657 |
|  | Acidosis | 53 [37–59] | 47 [39–55] | 49 [31–52] | 44 [28–50] |  |  |  |
| RAP (mm Hg) | Normal | 4 [2–6] | 3 [2–5] | 2 [2–5] | 2 [2–5] | <0.001 | 0.200 | 0.298 |
|  | Acidosis | 4 [2–5] | 5 [4–6] | 4 [3–5] | 4 [2–4] |  |  |  |
| SVRI (dynes•sec•cm<sup>-5</sup>/kg) | Normal | 287.7 [219.2–326.2] | 247.2 [134.0–332.9] | 154.5 [93.9–213.3]<sup>A</sup> | 100.6 [65.8–123.9]<sup>AB</sup> | <0.001 | <0.001 | 0.003 |
|  | Acidosis | 140.5 [113.7–248.0]* | 132.4 [110.5–238.3] | 99.7 [90.8–155.7] | 80.3 [67.9–133.0] |  |  |  |
| PAP (mm Hg) | Normal | 11 [10–12] | 11 [8–13] | 12 [8–16] | 13 [10–18] | <0.001 | 0.167 | 0.948 |
|  | Acidosis | 19 [17–21]* | 19 [18–19]* | 20 [18–21]* | 20 [19–22]* |  |  |  |
| PAOP (mm Hg) | Normal | 4 [4–6] | 4 [3–5] | 3 [3–5] | 3 [2–6] | <0.001 | <0.001 | 0.454 |
|  | Acidosis | 11 [9–12]* | 10 [6–14]* | 9 [7–11]* | 9 [7–9]*<sup>A</sup> |  |  |  |
| PVRI (dynes•sec•cm<sup>-5</sup>/kg) | Normal | 24.6 [16.9–49.9] | 19.7 [16.4–37.6] | 19.0 [11.5–33.7] | 19.8 [8.6–25.8] | 0.661 | 0.811 | 0.201 |
|  | Acidosis | 19.2 [12.6–22.4] | 20.4 [11.5–34.2] | 22.7 [13.2–31.3] | 21.4 [14.1–23.7] |  |  |  |

CI, cardiac index; HR, heart rate; SVI, stroke volume index; SAP, systolic arterial pressure; MAP, mean arterial pressure; DAP, diastolic arterial pressure; RAP, right atrial pressure; SVRI, systemic vascular resistance index; PAP, pulmonary arterial pressure; PAOP, pulmonary arterial occlusion pressure; PVRI, pulmonary vascular resistance index. The rank transformation version of two-way ANOVA applies on condition and dobutamine treatment. Superscript A, B, and C show significant difference ( $P < 0.05$ ) from baseline, 0.3 µg/kg/min, and 0.6 µg/kg/min by Steel-Dwass, respectively. \* shows a significant difference ( $P < 0.05$ ) as compared with each dose in Normal condition.

CI and HR significantly (*P* < 0.001) increased in response to colforsin administration relative to the baseline. In contrast, compared to values at baseline, colforsin significantly lowered SAP (*P* < 0.05), MAP (*P* < 0.05), SVRI (*P* < 0.001), and PAOP (*P* < 0.001). The numbers of dogs with mean PAP > 20 mm Hg were one (17%) at baseline, zero (0%) at 0.3 μg/kg/min, three (50%) at 0.6 μg/kg/min, and four (67%) at 1.2 μg/kg/min in the Acidosis condition.

Miosis was observed in three dogs (50%) receiving colforsin and four (67%) in the Acidosis condition but it disappeared within 6 h after the end of colforsin infusion. Nausea or vomiting was transiently observed in one dog (17%) in the Normal condition and two dogs (33%) in the Acidosis condition. All dogs ate and drank within 3 h after the end of the experiment. No dog presented with complications as a result of the drug administration according to their blood chemistry and general physical examinations 2 weeks after the end of the experiment.

### Blood gases and biochemical analyses in response to dobutamine

The blood gas and biochemical test results at baseline and at the end of the experiment are shown in Table 3. The pH at baseline was 7.38 (range 7.33–7.41) in the Normal condition and 6.99 (range 6.96–7.05) in the Acidosis condition. In both cases, the pH had slightly decreased by the end of the experiment. The baseline PaCO_2_ was 38.2 mm Hg (range 36.0–42.3 mm Hg) in the Normal condition and 109.0 mm Hg (range 101.0–114.3 mm Hg) in the Acidosis condition. Relative to the baseline, arterial oxygen delivery (DaO_2_) was significantly increased by dobutamine administration in both conditions (*P* < 0.05).

**Table 3.** The effects of dobutamine on blood gas examination and blood biochemical test in six anesthetized dogs in eucapnia (Normal) and acute respiratory acidosis (Acidosis) at baseline and at the end of the experiment.

| Variable (Unit) | Condition | Baseline | End of experiment | Reference |
| --- | --- | --- | --- | --- |
| Adrenaline (ng/mL) | Normal | 0.01 [0.01–0.13] | — | <0.10 |
|  | Acidosis | 0.26 [0.08–2.08]§ | — |  |
| Noradrenaline (ng/mL) | Normal | 0.04 [0.02–0.09] | — | 0.10–0.50 |
|  | Acidosis | 0.32 [0.24–0.44]§ | — |  |
| Dopamine (ng/mL) | Normal | 0.01 [0.01–0.02] | — | <0.03 |
|  | Acidosis | 0.02 [0.01–0.03] | — |  |
| PCV (%) | Normal | 34 [27–39] | 35 [33–43] | 37–55 |
|  | Acidosis | 40 [34–49] | 45 [40–51] |  |
| pH | Normal | 7.38 [7.33–7.41] | 7.30 [7.25–7.36]* | 7.35–7.45 |
|  | Acidosis | 6.99 [6.96–7.05]† | 6.92 [6.86–6.95]‡ |  |
| PaCO<sub>2</sub> (mm Hg) | Normal | 38 [36–42] | 42 [35–46] | 30.8–42.8 |
|  | Acidosis | 109 [101–114]† | 126 [115–146]‡ |  |
| PaO<sub>2</sub> (mm Hg) | Normal | 539 [495–571] | 579 [568–607]* | 80.9–103.3 |
|  | Acidosis | 525 [443–551] | 505 [473–544] |  |
| HCO<sub>3</sub><sup>-</sup> (mEq/L) | Normal | 22.8 [21.9–24.6] | 20.1 [18–21.8]* | 18.8–25.6 |
|  | Acidosis | 26.6 [24.6–27.9]† | 25.9 [24.7–27.9] |  |
| DaO<sub>2</sub>I (mL O<sub>2</sub>/min/kg) | Normal | 31.7 [20.6–40.7] | 89.9 [67.4–109.6]* | — |
|  | Acidosis | 49.1 [44.5–59.7]† | 101.1 [83.3–112.4]‡ |  |
| VaO<sub>2</sub>I (mL O<sub>2</sub>/min/kg) | Normal | 4.0 [2.4–4.3] | 7.0 [5.5–7.8]* | — |
|  | Acidosis | 4.4 [3.3–6.2] | 6.5 [4.2–7.3] |  |
| O<sub>2</sub>ER (%) | Normal | 13.5 [11.2–17.0] | 7.8 [6.1–9.5]* | — |
|  | Acidosis | 8.4 [7.1–11.9] | 6.1 [4.6–7.9] |  |
| BEecf (mEq/L) | Normal | -2.0 [-3.2–-1.0] | -7.0 [-8.1–-4.0] | -4–+4 |
|  | Acidosis | -5.0 [-7.0–-2.6] | -6.6 [-8.0–-5.2] |  |
| Lactate (mmol/L) | Normal | 1.5 [0.6–3.2] | 0.3 [0.3–0.5]* | <2.0 |
|  | Acidosis | 0.6 [0.3–0.9] | 0.6 [0.3–1.6] |  |
| Na (mEq/L) | Normal | 144 [142–148] | 146 [143–148] | 135–147 |
|  | Acidosis | 146 [144–148] | 146 [141–148] |  |
| K (mEq/L) | Normal | 3.7 [3.3–4.3] | 3.5 [3.0–4.8] | 3.5–5.0 |
|  | Acidosis | 3.8 [3.2–4.3] | 6.1 [5.0–7.5]‡ |  |
| Cl (mEq/L) | Normal | 115 [111–119] | 116 [115–123] | 95–125 |
|  | Acidosis | 113 [109–118] | 113 [111–117] |  |
| Glucose (mg/dL) | Normal | 103 [89–137] | 94 [87–106] | 60–110 |
|  | Acidosis | 150 [123–200]† | 138 [121–243] |  |
| BUN (mg/dL) | Normal | 15.0 [12.1–17.4] | 13.0 [11.0–15.7] | 10–20 |
|  | Acidosis | 15.0 [11.0–22.0] | 18.5 [15.0–25.0] |  |
| Creatinin (mg/dL) | Normal | 0.5 [0.4–0.9] | 0.35 [0.3–0.5] | 0.6–1.2 |
|  | Acidosis | 0.6 [0.5–0.7] | 1.05 [0.6–1.4] |  |

PCV, packed cell volume; PaCO<sub>2</sub>, arterial partial pressure of carbon dioxide; PaO<sub>2</sub>, arterial partial pressure of oxygen; HCO<sub>3</sub><sup>-</sup>, bicarbonate ion; DaO<sub>2</sub>I, oxygen delivery; VaO<sub>2</sub>I, oxygen consumption; O<sub>2</sub>ER, oxygen extraction ratio; BEecf, base excess in the extracellular fluid; Na, sodium ion; K, potassium ion; Cl, chloride ion; BUN, blood urea nitrogen. The reference values were shown from individual testing apparatus. § shows significant difference ( $P < 0.05$ ) from baseline in Normal condition by Wilcoxon signed-rank test. \* and † show significant difference ( $P < 0.05$ ) from baseline in Normal condition, respectively, and ‡ shows significant difference ( $P < 0.05$ ) from baseline in Acidosis condition by Steel-Dwass, respectively.

Blood adrenaline and noradrenaline levels were significantly higher in the Acidosis condition than the Normal condition at baseline (*P* < 0.05). Plasma glucose was significantly (*P* < 0.05) higher in the Acidosis condition than the Normal condition. In the latter case, plasma potassium at the end of the experiment was significantly (*P* < 0.05) higher than it was at baseline.

### Cardiovascular effects of dobutamine

The effects of dobutamine on the cardiovascular parameters in the Normal and Acidosis conditions are shown in Table 4. There were no interactions between dobutamine treatment and pH in terms of their effects on the cardiovascular parameters. The CI, SVI, PAP, and PAOP were higher and the SVRI was lower in the Acidosis condition than the Normal condition at baseline, and the differences were significant (*P* < 0.05). Relative to the baseline value, the rate of increase in CI in the Acidosis condition was greater than that in the Normal condition (Normal vs. Acidosis: 46% vs. 44%, 5 μg/kg/min; 129% vs. 66%, 10 μg/kg/min; 157% vs. 82%, 20 μg/kg/min). The CI, HR, SVI, and PAP significantly increased in response to dobutamine administration (*P* < 0.001). Dobutamine administration significantly lowered SAP (*P* < 0.01), MAP (*P* < 0.01), DAP (*P* < 0.001), and SVRI (*P* < 0.001) compared to levels at baseline. The numbers of dogs with mean PAP > 20 mm Hg were one (17%) at 10 μg/kg/min and three (50%) at 20 μg/kg/min in the Normal condition, two (33%) at baseline, and six (100%) > 5 μg/kg/min in the Acidosis condition.

**Table 4.**
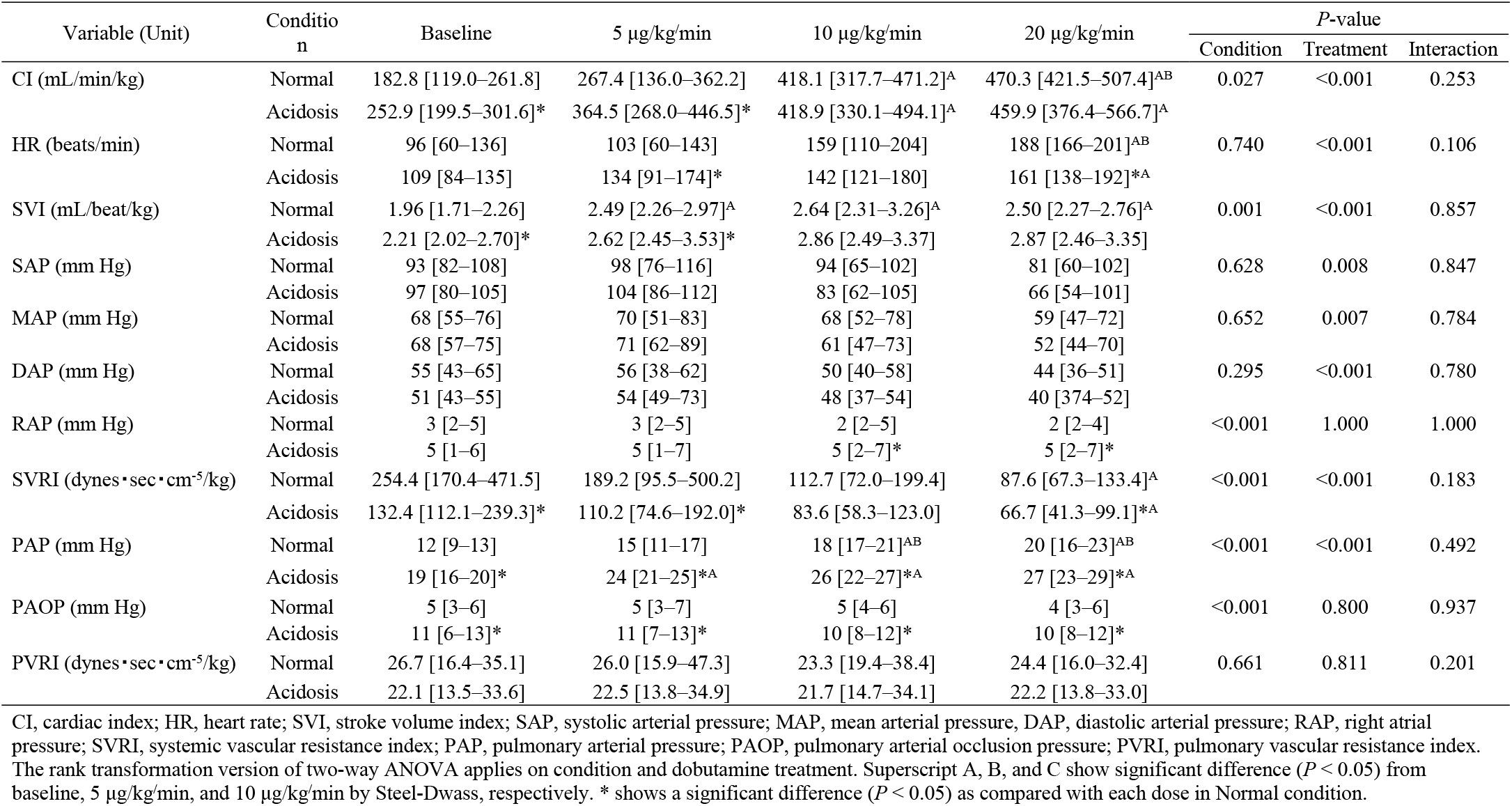
Median [range] values for cardiovascular variables at baseline, 5 μg/kg/min, 10 μg/kg/min, and 20 μg/kg/min dose of intravenous dobutamine in six anesthetized dogs in eucapnia (Normal) and acute respiratory acidosis (Acidosis) conditions.

| Variable (Unit) | Condition | Baseline | 5 µg/kg/min | 10 µg/kg/min | 20 µg/kg/min | P-value |  |  |
| --- | --- | --- | --- | --- | --- | --- | --- | --- |
|  |  |  |  |  |  | Condition | Treatment | Interaction |
| CI (mL/min/kg) | Normal | 182.8 [119.0–261.8] | 267.4 [136.0–362.2] | 418.1 [317.7–471.2]<sup>A</sup> | 470.3 [421.5–507.4]<sup>AB</sup> | 0.027 | <0.001 | 0.253 |
|  | Acidosis | 252.9 [199.5–301.6]* | 364.5 [268.0–446.5]* | 418.9 [330.1–494.1]<sup>A</sup> | 459.9 [376.4–566.7]<sup>A</sup> |  |  |  |
| HR (beats/min) | Normal | 96 [60–136] | 103 [60–143] | 159 [110–204] | 188 [166–201]<sup>AB</sup> | 0.740 | <0.001 | 0.106 |
|  | Acidosis | 109 [84–135] | 134 [91–174]* | 142 [121–180] | 161 [138–192]*<sup>A</sup> |  |  |  |
| SVI (mL/beat/kg) | Normal | 1.96 [1.71–2.26] | 2.49 [2.26–2.97]<sup>A</sup> | 2.64 [2.31–3.26]<sup>A</sup> | 2.50 [2.27–2.76]<sup>A</sup> | 0.001 | <0.001 | 0.857 |
|  | Acidosis | 2.21 [2.02–2.70]* | 2.62 [2.45–3.53]* | 2.86 [2.49–3.37] | 2.87 [2.46–3.35] |  |  |  |
| SAP (mm Hg) | Normal | 93 [82–108] | 98 [76–116] | 94 [65–102] | 81 [60–102] | 0.628 | 0.008 | 0.847 |
|  | Acidosis | 97 [80–105] | 104 [86–112] | 83 [62–105] | 66 [54–101] |  |  |  |
| MAP (mm Hg) | Normal | 68 [55–76] | 70 [51–83] | 68 [52–78] | 59 [47–72] | 0.652 | 0.007 | 0.784 |
|  | Acidosis | 68 [57–75] | 71 [62–89] | 61 [47–73] | 52 [44–70] |  |  |  |
| DAP (mm Hg) | Normal | 55 [43–65] | 56 [38–62] | 50 [40–58] | 44 [36–51] | 0.295 | <0.001 | 0.780 |
|  | Acidosis | 51 [43–55] | 54 [49–73] | 48 [37–54] | 40 [37–52] |  |  |  |
| RAP (mm Hg) | Normal | 3 [2–5] | 3 [2–5] | 2 [2–5] | 2 [2–4] | <0.001 | 1.000 | 1.000 |
|  | Acidosis | 5 [1–6] | 5 [1–7] | 5 [2–7]* | 5 [2–7]* |  |  |  |
| SVRI (dynes•sec•cm<sup>-5</sup>/kg) | Normal | 254.4 [170.4–471.5] | 189.2 [95.5–500.2] | 112.7 [72.0–199.4] | 87.6 [67.3–133.4]<sup>A</sup> | <0.001 | <0.001 | 0.183 |
|  | Acidosis | 132.4 [112.1–239.3]* | 110.2 [74.6–192.0]* | 83.6 [58.3–123.0] | 66.7 [41.3–99.1]*<sup>A</sup> |  |  |  |
| PAP (mm Hg) | Normal | 12 [9–13] | 15 [11–17] | 18 [17–21]<sup>AB</sup> | 20 [16–23]<sup>AB</sup> | <0.001 | <0.001 | 0.492 |
|  | Acidosis | 19 [16–20]* | 24 [21–25]*<sup>A</sup> | 26 [22–27]*<sup>A</sup> | 27 [23–29]*<sup>A</sup> |  |  |  |
| PAOP (mm Hg) | Normal | 5 [3–6] | 5 [3–7] | 5 [4–6] | 4 [3–6] | <0.001 | 0.800 | 0.937 |
|  | Acidosis | 11 [6–13]* | 11 [7–13]* | 10 [8–12]* | 10 [8–12]* |  |  |  |
| PVRI (dynes•sec•cm<sup>-5</sup>/kg) | Normal | 26.7 [16.4–35.1] | 26.0 [15.9–47.3] | 23.3 [19.4–38.4] | 24.4 [16.0–32.4] | 0.661 | 0.811 | 0.201 |
|  | Acidosis | 22.1 [13.5–33.6] | 22.5 [13.8–34.9] | 21.7 [14.7–34.1] | 22.2 [13.8–33.0] |  |  |  |

CI, cardiac index; HR, heart rate; SVI, stroke volume index; SAP, systolic arterial pressure; MAP, mean arterial pressure; DAP, diastolic arterial pressure; RAP, right atrial pressure; SVRI, systemic vascular resistance index; PAP, pulmonary arterial pressure; PAOP, pulmonary arterial occlusion pressure; PVRI, pulmonary vascular resistance index. The rank transformation version of two-way ANOVA applies on condition and dobutamine treatment. Superscript A, B, and C show significant difference ( $P < 0.05$ ) from baseline, 5 µg/kg/min, and 10 µg/kg/min by Steel-Dwass, respectively. \* shows a significant difference ( $P < 0.05$ ) as compared with each dose in Normal condition.

Atrial stasis was observed by the end of the experiment in one acidotic dog receiving dobutamine. Its plasma potassium was 7.5 mmol/L. After the experiment, its cardiac rhythm reverted to a normal electrocardiogram waveform. No other arrhythmia was observed. Miosis was observed in four dogs (67%) receiving dobutamine in the Acidosis condition. However, the miosis disappeared within 6 h after the end of dobutamine infusion. Nausea or vomiting was transiently observed in three dogs (50%) in the Normal condition and two dogs (33%) in the Acidosis condition. All dogs ate and drank within 3 h after the end of the experiment. No dog presented with complications as a result of the drug administration according to their blood chemistry and general physical examinations 2 weeks after the end of the experiment.

## Discussion

To the best of our knowledge, this study is the first to evaluate the dose-dependent cardiovascular function of colforsin in dogs. Colforsin had a cardiovascular action similar to that of dobutamine. It increased CI and HR and decreased SVRI in a dose-dependent manner. However, under acute respiratory acidosis, the rates of change in CI, HR, and SVRI were attenuated with both colforsin and dobutamine. Therefore, colforsin and dobutamine doses may have to be increased under respiratory acidosis.

### 1) Cardiovascular effects of dobutamine and colforsin in the Normal condition

Dobutamine is a synthetic dopamine analog which stimulates beta-1, beta-2, and alpha-1 adrenoceptors in the cardiovascular system at doses approximating those used clinically (1–20 μg/kg/min) [2,9,10]. The inotropic activity of dobutamine is the result of stimulating both beta-1 and alpha-1 adrenoceptors in the myocardium. Furthermore, the beta-2 adrenoceptor-mediated vasodilatory effect of dobutamine is offset by alpha-1 adrenoceptor-mediated vasoconstrictor activity. Therefore, dobutamine increases CI and HR and decreases SVRI (inodilation) in a dose-dependent manner [11,12]. In the present study, dobutamine administration raised both CI and HR and lowered SVRI which corroborates previous reports.

Colforsin activates adenylate cyclase in cardiomyocytes and vascular smooth muscle without mediating catecholamine beta adrenoceptors. It increases cardiac contractility and reduces peripheral vascular resistance [13]. We used dobutamine as a positive control in the present study. At clinical doses, dobutamine induced dose-related inotropism and afterload reduction with a relative lack of chronotropism. These conditions are appropriate for the management of patients with congestive heart failure. They could also improve renal blood flow by enhancing cardiac output and beta-2 adrenoceptor-stimulated vasodilation [12,14,15]. The colforsin doses administered in the present study (0.3 μg/kg/min, 0.6 μg/kg/min, and 1.2 μg/kg/min) were those required to increase the heart rate to a level equivalent to that induced by dobutamine in our preliminary study (data not shown). In the present study, colforsin administration increased CI and HR and decreased SVRI as did dobutamine. Therefore, colforsin could substitute for dobutamine as an inodilator and might be useful for the treatment of pathological conditions such as congestive heart failure.

Unlike dobutamine, colforsin did not increase the PAP. Vascular smooth muscle in the pulmonary artery was relaxed by beta adrenoceptor stimulation [16]. In terms of inodilator dose–response effects in rats, dobutamine increased systolic pulmonary artery pressure [17]. It also slightly elevated pulmonary vascular resistance in anesthetized dog [18]. Pulmonary hypertension is defined as pulmonary arterial systolic pressure > 30 mm Hg or pulmonary arterial mean pressure > 20 mm Hg [19]. In the present study, dobutamine increased the PAP in a dose-dependent manner. Dobutamine at 10 μg/kg/min elevated the mean PAP > 20 mm Hg in 1/6 dogs while 20 μg/kg/min dobutamine had the same effect on 3/6 dogs. In contrast, colforsin administration produced no pulmonary hypertension. Left-sided heart disease is the most common cause of pulmonary hypertension in humans and dogs [19,20]. In a study of 60 dogs with pulmonary hypertension, 38 (63%) presented with degenerative mitral valve disease [21]. Other studies indicated that 14–31% of all dogs diagnosed with the latter disorder developed pulmonary hypertension [22,23]. Therefore, colforsin might be more efficacious than dobutamine in the treatment of severe mitral valve insufficiency accompanied by pulmonary hypertension. The effects of colforsin and dobutamine on the vascular smooth muscle of the pulmonary artery merit further investigation.

### 2) Cardiovascular effects of dobutamine and colforsin in the Acidosis condition

In the present study, the acute respiratory acidosis canine model was induced by carbon dioxide inhalation [24]. The baseline pH at the time of dobutamine and colforsin administration was ~7.0. However, PaCO_2_ slightly increased during the experiment (0.5 h for stabilization and 3 h for measurement). Although the pH had slightly decreased by the end of the experiment, we believe that acute respiratory acidosis was induced in the dogs at ~pH 7.0.

Acute respiratory acidosis increases cardiac output and heart rate in dogs [25]. Symptoms of early hypercapnia include nausea/vomiting, muscle twitching, extrasystoles, and sympathetic nervous system stimulation. In the present study, the plasma adrenaline and noradrenaline levels in the Acidosis condition were significantly higher than those in the Normal condition. Therefore, elevated catecholamines could increase cardiac output and stroke volume under acidosis. Hypercapnia also causes anesthesia and peripheral blood vessel dilation [25]. At baseline, hypercapnia might have decreased systemic vascular resistance under acidosis in the present study.

The affinity of catecholamine for the beta adrenoceptor decreases under acidosis [26]. Therefore, the cardiovascular effects of catecholamines are attenuated. On the other hand, colforsin improved cardiac contractility in isolated and acid-perfused rat heart under acidosis [7]. It was also reported that the cAMP level in cardiac muscle cells was higher in response to colforsin than to catecholamines [7]. However, the cardiovascular effects of colforsin on CI, HR, and SVRI resembled those of dobutamine in the present study. The rates of change in these variables in response to both drugs were weaker under the Acidosis condition than under the Normal condition. As cardiomyocyte cAMP was not measured here, it could not be determined whether the same reaction was occurring as that reported in the previous study. Moreover, acidic perfusate rather than hypercapnia was considered in that study. Hypercapnia is anesthetic and suppresses cardiovascular function [27,28]. Consequently, even if the same isoflurane dose was administered to both groups, anesthesia may have been more profound in the Acidosis group than in the Normal group because the former presented with hypercapnia. For this reason, the effects of colforsin and dobutamine may have been attenuated by hypercapnia.

Pulmonary arterial vasoconstriction occurs in response to alpha adrenoceptor the stimulation [29]. The alpha adrenoceptors in the pulmonary arteries have a high affinity for catecholamines such as noradrenaline. As the baseline noradrenaline concentration was high in the present study, the PAP was higher in the Acidosis condition than in the Normal condition. In the experimental induction of microembolic pulmonary hypertension, high dobutamine doses decreased pulmonary artery pressure [30]. In the present study, all dogs administered with dobutamine showed pulmonary hypertension > 20 mm Hg. Although certain dogs presented with pulmonary hypertension at high colforsin doses, their PAP was low relative to that induced by dobutamine. Even under acidosis, the influence of colforsin on pulmonary artery pressure was small compared with that of dobutamine. Therefore, the fact that colforsin had zero impact on pulmonary artery pressure may facilitate its application as an adjunct to (or replacement for) dobutamine.

Acute respiratory distress syndrome (ARDS) is life-threatening and caused by sepsis or a systemic inflammatory response. ARDS requires ventilator management in intensive care and lung protective ventilation is recommended [31]. A consequence of low tidal volume ventilation is an elevation in PaCO_2_. In humans, high PaCO_2_ levels (> 70 mm Hg) may be tolerated (permissive hypercapnia). Nevertheless, heavier sedation or paralysis may be required to prevent patient-ventilator asynchrony [32,33]. Past evidence from experimental animal studies [24,34] and human clinical trials [31] suggest that lung-protective ventilation would also be warranted in veterinary patients. We set our acute respiratory acidosis model higher than that required for lung-protective ventilation in order to differentiate the drug effects clearly. ARDS often causes pulmonary hypertension as a result of hypoxic pulmonary vasoconstriction and pulmonary blood vessel organization. In the present study, colforsin did not raise pulmonary artery pressure in the Acidosis condition. It was reported that colforsin attenuates bronchoconstriction- and pulmonary hypertension-induced serotonin infusion in dogs [35]. Therefore, colforsin may be able to improve cardiac function in permissive hypercapnia with pulmonary hypertension more effectively than dobutamine. The application of colforsin for the treatment of pulmonary hypertension caused by mitral valve insufficiency and ARDS might be a new therapeutic strategy in both veterinary and human medicine.

In the present study, plasma glucose level was higher in the Acidosis condition than in the Normal condition at baseline. Catecholamines markedly increase plasma glucose levels [36,37]. Insulin secretion declines after alpha adrenoceptor activation but rises in response to beta-2 adrenoceptor activation [38]. The high baseline plasma glucose level in the Acidosis condition was positively correlated with high plasma adrenaline and noradrenaline levels. Although dobutamine stimulates beta-1, beta-2, and alpha-1 adrenoceptors [2,9], the dobutamine dosage administered in this present study did not affect plasma glucose level under the Normal condition. The effects of alpha-1 catecholamine may have been offset by the beta-2 catecholaminic action of dobutamine. The effects of colforsin on insulin secretion are unknown. Nevertheless, the colforsin dose administered in the present study had no effect on the plasma glucose level. Therefore, colforsin might be appropriate for diabetic patients whose cardiovascular function must be improved without raising their plasma glucose levels. In the future, the influence of colforsin administration on plasma insulin concentration should be investigated.

In the Acidosis condition, the baseline plasma potassium level was higher than that at the end of the experiment. Lactic acidosis is probably not associated with major intracellular shifts in potassium level. However, respiratory acidosis may influence the serum potassium concentration [39]. In the Normal condition, neither dobutamine nor colforsin increased plasma potassium levels. Moreover, there was no significant difference in plasma potassium between the Normal and Acidosis conditions at baseline. Relative to the baseline, however, plasma potassium level was significantly higher in the Acidosis condition at the end of experiment. Plasma potassium level rose in 3.5 h (0.5 h for stabilization and 3 h for the experiment) after the induction of acute respiratory acidosis. One acidotic dog receiving dobutamine (potassium level = 7.5 mmol/L) showed atrial stasis at the end of the experiment. Since the sample size was small in this assay, we could not confirm the relationship between dobutamine and arrhythmia. On the other hand, no arrhythmia was observed in dogs receiving colforsin (maximum potassium level = 8.3 mmol/L). Although there was no atrial stasis, colforsin acted as an inodilator here. Colforsin also suppressed digitalis- and epinephrine-induced ventricular arrhythmia models in dogs [40]. Abnormal plasma potassium levels and arrhythmia are often observed in heart- and renal failure [41]. Neither colforsin nor dobutamine affected plasma potassium levels under eucapnia. Therefore, both drugs neither aggravate nor alleviate acidotic increases in plasma potassium. For these reasons, colforsin could substitute for dobutamine in heart- and renal failure therapy. In the future, the associations among colforsin, plasma potassium level, and arrhythmia in these diseases should be investigated.

Some dogs presented with transient nausea or vomiting during recovery from both drugs. Dobutamine has provoked nausea, headache, vomiting, and dyspnea [2]. To the best of our knowledge, adverse effects have not been reported for colforsin. Nevertheless, it still may have side effects similar to those of dobutamine. The aim of the present study was to investigate the cardiovascular effects of colforsin under acidosis. Since the dose administered was impractical, many side effects may have been induced. In future research, we could endeavor to optimize the dosage of colforsin which would improve cardiovascular function in the presence of respiratory- or other acidosis. Certain dogs under the Acidosis condition showed miosis at the end of the experiment. Miosis occurs when intracranial pressure increases. Since hypercapnia increases cerebral blood flow [42], it may have also elevated intracranial pressure and induced miosis. Dogs presenting with miosis returned to normal pupillary diameter within 6 h after the experiment. No other neurological complications were observed. Although acute respiratory acidosis was maintained for 3.5 h in the present study, pupil size should be verified in respiratory acidosis and permissive hypercapnia in a clinical setting.

We conducted this study assuming that pulmonary edema or ARDS may complicate respiratory acidosis. However, there were certain limitations here. Although oxygenation is impaired in pulmonary edema and ARDS, we did not conduct this experiment under hypoxemia which stimulates the sympathetic nervous system and enhances cardiovascular function. We wanted to clarify the cardiovascular effects of colforsin in normal dogs. Therefore, we conducted this experiment with 100% oxygen carrier gas. Next, we used healthy dogs free of heart or lung disease. Dogs with severe mitral valve insufficiency causing pulmonary edema or ARDS already have depressed cardiorespiratory function. Therefore, administering colforsin and dobutamine to these patients may produce different cardiovascular effects. Further studies are needed to establish the effects of colforsin and dobutamine on these disease models and clinical cases and to verify the safety and efficacy of colforsin. In turn, these findings could be adapted to human medicine.

## Conclusions

The cardiovascular effects of colforsin and dobutamine are similar in healthy beagles under isoflurane anesthesia. In acute respiratory acidosis induced by carbon dioxide inhalation, cardiovascular function was enhanced by endogenous catecholamine secretion. In addition, the rates of change in CI, HR, and SVRI caused by colforsin and dobutamine administration were attenuated. Therefore, it may be necessary to increase the colforsin and dobutamine doses under respiratory acidosis relative to those administered under the normal condition. Since colforsin had little effect on the PAP, it may be more suitable as an inodilator than dobutamine in the treatment of diseases which increase the PAP. Our next steps are to induce a pulmonary hypertension canine model, confirm the effects of colforsin on it, adapt colforsin administration for patients with pulmonary hypertension in our institution, and compare its efficacy with that of existing catecholamines.

## Acknowledgments

The authors thank Dr. Kohei Makita for guidance in the use of the statistics software package R and Editage (www.editage.jp) for English language editing.

The role of each author is as follows:

| Name | Role |
| --- | --- |
| Takaharu Itami | Conceptualization, data curation, formal analysis, investigation, and original draft preparation |
| Kiwamu Hanazono | Conceptualization, data curation, formal analysis, investigation, and methodology |
| Norihiko Oyama | Investigation |
| Tadashi Sano | Investigation |
| Kazuto Yamashita | Investigation and supervision |

